# The anti-dyskinetic and the anti-parkinsonian effects of sub-anesthetic ketamine are also present in female preclinical models

**DOI:** 10.64898/2026.10.27.762342

**Authors:** Raveena Parmar, Mitchell J. Bartlett, Jasmine R. Meredith, Sansita Singh, Sydney Ragsdale, Scott J. Sherman, Torsten Falk

## Abstract

Biological sex is an important determinant of neuropharmacological response, yet females remain underrepresented in preclinical studies of Parkinson’s disease (PD) and L-DOPA-induced dyskinesia (LID). We previously demonstrated that sub-anesthetic ketamine reduces established LID, attenuates dyskinesia development, and produces anti-parkinsonian effects in male unilateral 6-hydroxydopamine (6-OHDA) rat models. However, whether these therapeutic effects extend to females remained unknown. Here, we evaluated the anti-dyskinetic and anti-parkinsonian effects of sub-anesthetic ketamine in female Sprague-Dawley rats using the attenuation and established models of LID. In the attenuation paradigm, unilateral 6-OHDA-lesioned females received repeated L-DOPA administration with either ketamine (20 mg/kg, i.p.; 10-h dosing paradigm) or vehicle. Ketamine significantly attenuated the progressive development of dyskinesia, indicating that early intervention modified the trajectory of LID expression. In the established LID paradigm, ketamine produced robust significant acute reductions in abnormal involuntary movements (AIMs) after dyskinesia had fully developed, and these significant effects persisted for up to 49 days following treatment, demonstrating sustained suppression of dyskinesia severity. To determine whether ketamine altered therapeutic motor function, forelimb adjusting steps testing was performed in female unilateral 6-OHDA rats. Ketamine did not impair L-DOPA-mediated motor improvement suggesting compatibility with the gold standard medication. Ketamine was also by itself associated with significantly improved stepping performance during ketamine treatment conditions, indicating acute anti-parkinsonian motor benefit. Together, these findings demonstrate that the anti-dyskinetic and anti-parkinsonian effects of ketamine extend to female preclinical models and strengthen the translational rationale for repurposing sub-anesthetic ketamine as a potential therapeutic strategy for individuals with LID.

**Highlights:**

- Ketamine attenuated development of L-DOPA-induced dyskinesia in females
- Ketamine reduced established dyskinesia acutely and had long term effects
- Ketamine preserved L-DOPA-mediated anti-parkinsonian effects
- Ketamine alone improved forelimb stepping in female 6-OHDA rats
- Therapeutic efficacy of ketamine for Parkinson’s disease extends to female models

## Introduction

Despite growing recognition of biological sex as an important determinant of neuropharmacological responses, females remain underrepresented in preclinical studies of Parkinson’s disease (PD), L-DOPA-induced dyskinesia (LID), and anti-dyskinetic therapies.^1,2^ Although sub-anesthetic ketamine has demonstrated anti-dyskinetic efficacy in male rodent models, whether these effects are also observed in females has not been established.^3,4^ This leaves an important gap in the preclinical evaluation of ketamine as a potential therapeutic for LID, particularly given evidence that sex influences dopaminergic neurotransmission, synaptic plasticity, and corticostriatal circuit function, processes that contribute to the development and expression of dyskinesia and may influence therapeutic responsiveness.^5,6^

Clinical and experimental evidence further supports the relevance of biological sex to PD and LID. Sex differences have been reported in PD clinical presentation, dopaminergic responsiveness, and susceptibility to L-DOPA-associated motor complications.^5,6,7^ Sex-dependent modulation of striatal dopamine signaling may be particularly relevant to LID, as gonadal hormones, including estrogens, can influence dopamine synthesis, release, receptor function, and neuronal responsiveness.^8,9^ Because LID arises in part from maladaptive plasticity within dopamine-and glutamate-regulated corticostriatal circuits, sex-related differences in these processes may contribute to variability in both dyskinesia development and therapeutic response.

Dysregulated glutamatergic neurotransmission and N-methyl-D-aspartate (NMDA) receptor signaling are strongly implicated in the pathophysiology of LID. Chronic L-DOPA exposure induces maladaptive corticostriatal plasticity characterized by altered glutamatergic transmission, receptor regulation, and synaptic homeostasis.^3,10^ Picconi et al. demonstrated that dyskinetic states are associated with impaired bidirectional synaptic plasticity within the striatum, including abnormalities in both long-term potentiation and long-term depression.^11^ Consistent with these findings, Gardoni et al. identified alterations in NMDA receptor composition and receptor-scaffolding interactions within dyskinetic striatal circuitry.^12^ Collectively, these findings implicate disrupted glutamatergic signaling and NMDA receptor-dependent plasticity in LID and provide a mechanistic rationale for pharmacological approaches targeting these pathways.

Ketamine is a non-competitive NMDA receptor antagonist that modulates glutamatergic neurotransmission and synaptic plasticity.^13^ At sub-anesthetic doses, ketamine produces rapid and sustained neurobiological effects involving activity-dependent brain-derived neurotrophic factor (BDNF) signaling, enhanced alpha-amino-3-hydroxy-5-methyl-4-isoxazolepropionic acid (AMPA) receptor-mediated signaling, and downstream pathways associated with synaptic remodeling.^15,16,17,18^ Although these mechanisms have been characterized predominantly in neuropsychiatric models, their capacity to modify maladaptive plasticity provides a plausible mechanistic basis for ketamine’s effects on LID-associated circuit dysfunction. Consistent with this rationale, previous work from our laboratory demonstrated that sub-anesthetic ketamine attenuates both the development and established expression of LID and produces anti-parkinsonian effects in male rodent models.^4,19,20^

Sex-dependent differences in ketamine responsiveness have been demonstrated in preclinical models of depression. Although depression and LID are clinically distinct disorders involving different neural circuits, ketamine engages neurobiological processes relevant to both conditions, including glutamatergic signaling and activity-dependent plasticity. Findings from depression models may therefore inform our understanding of how biological sex influences ketamine responsiveness in LID. Carrier and Kabbaj reported that female rats exhibited antidepressant-like responses to a low dose of ketamine that was ineffective in males, indicating greater sensitivity to ketamine in females.^21^ This enhanced sensitivity was abolished following ovariectomy and restored by estrogen and progesterone replacement, suggesting that ovarian hormones contribute to sex-dependent ketamine responsiveness.^21^ Subsequent work further demonstrated that behavioral and biochemical responses to low-dose ketamine can vary according to sex and estrous-cycle-associated hormonal status.^22^ Sex-dependent differences have also been reported in ketamine pharmacokinetics, including differences in plasma and brain exposure to ketamine and its metabolites following comparable dosing paradigms.^23^ Collectively, these findings indicate that biological sex can influence both the pharmacodynamic and pharmacokinetic properties of ketamine, providing a strong rationale for evaluating its therapeutic efficacy in female LID models.

The present study therefore builds upon previous findings in male LID models to determine whether the anti-dyskinetic and anti-parkinsonian effects of sub-anesthetic ketamine are also observed in female rats. Using paradigms designed to evaluate both the development and established expression of LID, we examined whether ketamine attenuates the emergence of dyskinesia when administered during early L-DOPA exposure and reduces dyskinetic behavior after stable abnormal involuntary movements (AIMs) have developed. Forepaw adjusting steps (FAS) were additionally assessed to evaluate the anti-parkinsonian effects of ketamine and determine whether ketamine altered the therapeutic motor benefit of L-DOPA. We hypothesized that sub-anesthetic ketamine would attenuate both developing and established LID while also improving parkinsonian motor deficits in female rats. Together, these experiments extend the preclinical evaluation of ketamine to females and determine whether its previously reported anti-dyskinetic and anti-parkinsonian effects generalize beyond male models.

## Material and Methods

**Table 1.** Abbreviations.

| **6-OHDA** | 6-Hydroxydopamine |
|---|---|
| **AADC** | Aromatic L-amino Acid Decarboxylase |
| **AIMs** | Abnormal Involuntary Movements |
| **AMPA** | α-Amino-3-hydroxy-5-methyl-4-isoxazolepropionic Acid |
| **AP** | Anterior–Posterior |
| **ARRIVE** | Animal Research: Reporting of *In Vivo* Experiments |
| **BCA** | Bicinchoninic Acid |
| **BDNF** | Brain-Derived Neurotrophic Factor |
| **CO<sub>2</sub>** | Carbon Dioxide |
| **DA** | Dopamine |
| **DMSO** | Dimethyl Sulfoxide |
| **DV** | Dorsal–Ventral |
| **FAS** | Forelimb Adjusting Steps |
| **i.p.** | Intraperitoneal |
| **LAO** | Limb, Axial, and Orolingual |
| **LID** | L-DOPA-Induced Dyskinesia |
| **L-DOPA** | Levodopa |
| **LTD** | Long-Term Depression |
| **LTP** | Long-Term Potentiation |
| **MFB** | Medial Forebrain Bundle |
| **ML** | Medial–Lateral |
| **mTOR** | Mechanistic Target of Rapamycin |
| **NIH** | National Institutes of Health |
| **NMDA** | N-Methyl-D-Aspartate |
| **PD** | Parkinson’s Disease |
| **RIPA** | Radioimmunoprecipitation Assay |
| **SEM** | Standard Error of the Mean |
| **TBST** | Tris-Buffered Saline with Tween-20 |
| **TH** | Tyrosine Hydroxylase |

### Animals

Adult female Sprague-Dawley rats (225 g at arrival; Envigo RMC Inc., Indianapolis, IN, USA) were housed two to three per cage in a temperature-and humidity-controlled room maintained on a 12-h reverse light/dark cycle. Food and water were provided ad libitum. All animal procedures were approved by the Institutional Animal Care and Use Committee at The University of Arizona and were conducted in accordance with the National Institutes of Health Guide for the Care and Use of Laboratory Animals and ARRIVE guidelines.

### Unilateral 6-hydroxydopamine (6-OHDA) lesion

Rats (n = 40; 250–300 g at time of surgery) were pretreated 30 min prior to 6-OHDA administration with desipramine hydrochloride (12.5 mg/kg, i.p.; Sigma-Aldrich, St. Louis, MO, USA) dissolved in sterile 0.9% saline containing 10% dimethyl sulfoxide to protect noradrenergic neurons. Animals were anesthetized with isoflurane (1.5–2.0%; VetOne, Boise, ID, USA) delivered in oxygen at 1.5 L/min using a vaporizer (JD Medical, Phoenix, AZ, USA) and secured in a stereotaxic frame.

6-OHDA hydrobromide (5.0 μg/μL in sterile 0.9% saline containing 0.02% ascorbic acid; Sigma-Aldrich) was infused into the right medial forebrain bundle at a rate of 0.5 μL/min using a microinjector (Stoelting Co., Wood Dale, IL, USA) attached to a 26-gauge needle and 10 μL Hamilton syringe (Hamilton Co., Reno, NV, USA). Two microliters of 6-OHDA were injected at each coordinate, corresponding to 10 μg of 6-OHDA per site. Injections were made at the following coordinates: AP = −1.8 mm, ML = +2.0 mm, DV = −8.2 mm and AP = −2.8 mm, ML = +1.8 mm, DV = −8.2 mm. The needle was left in place for 5 min following each infusion to minimize backflow before being slowly withdrawn. Consistent with established laboratory practice and the validated nature of the unilateral 6-OHDA lesion paradigm, sham-operated control groups were not included.^4,19,24^

### Amphetamine-induced rotation test

Lesion severity was assessed using amphetamine-induced rotational behavior. Rats received dextro-amphetamine sulfate (5 mg/kg, i.p.; Sigma-Aldrich) dissolved in sterile 0.9% saline and were placed individually in a plexiglass cylinder (38 cm diameter × 38 cm height). Ipsilateral and contralateral rotations were recorded for 1 min every 5 min over a 100-min testing period. Net ipsilateral rotations were calculated, and only animals exhibiting ≥4 net ipsilateral rotations/min were included in subsequent experiments. Animals meeting inclusion criteria were assigned to treatment groups balanced by lesion severity. Following lesion verification, 42 animals were included across the experiments. Experiment 1: female attenuation study (n = 20), and Experiment 2: female established study (n = 10), Experiment 3: female FAS study (n = 12).

### Drug and vehicle preparation

All pharmacological agents were administered intraperitoneally (i.p.) at a standardized injection volume of 1 mL/kg body weight. Ketamine (20 mg/kg; VetOne) and L-DOPA (6 or 7 mg/kg; Sigma-Aldrich) were prepared in 0.9% USP-grade sterile saline (VetOne). L-DOPA was co-administered with benserazide hydrochloride (14 mg/kg; Sigma-Aldrich), a peripheral aromatic L-amino acid decarboxylase inhibitor used to enhance central L-DOPA bioavailability and minimize peripheral dopamine conversion, consistent with established preclinical LID paradigms. Drug solutions were prepared fresh on the day of experimentation and administered according to treatment assignment. In Experiment 1 (“female attenuation study”), animals received L-DOPA (6 mg/kg, i.p.), whereas animals in Experiment 2 (“female established study”) received L-DOPA (7 mg/kg, i.p.) to maintain stable dyskinetic responding and to match the respective prior studies in males.^4,19^ Ketamine (20 mg/kg, i.p.) or vehicle was administered according to experimental design. All doses were calculated using individual body weight.

### Treatment groups and experimental design

#### Experiment 1: Female LID attenuation study

Experiment 1 evaluated whether sub-anesthetic ketamine attenuates the development of LID in female rats. Following 6-OHDA lesion verification, rats received daily L-DOPA (6 mg/kg, i.p.) with Benserazide (14 mg/kg, i.p.) for 14 consecutive days. AIMs were assessed every 3–4 days throughout the treatment period. On D0 and D7, animals received either ketamine (20 mg/kg, i.p.; n = 10) or vehicle (0.9% sterile saline; n = 10) using the full 10-h treatment paradigm. This consisted of five injections administered 2 h apart, with the fifth injection co-administered with L-DOPA. On non-treatment days, animals received L-DOPA without ketamine or vehicle pretreatment. Following the final behavioral assessment on D14, animals received a final ketamine or vehicle injection according to treatment assignment and were euthanized 1 h later for tissue collection.

#### Experiment 2: Female LID established study

Experiment 2 evaluated whether sub-anesthetic ketamine reduces established LID in female rats. Unilateral 6-OHDA-lesioned female rats received daily L-DOPA administration (7 mg/kg, i.p.) for 3 weeks to establish stable dyskinetic behavior. Following baseline establishment, animals continued receiving L-DOPA twice weekly (7 mg/kg, i.p.), and AIMs were reassessed every 4–7 days throughout the study. On D0, animals received ketamine treatment (*n* = 5) or vehicle (*n* = 5) using the 10-h dosing paradigm. Behavioral reassessments were subsequently conducted during continued L-DOPA exposure at D14, D18, D21, D28, D35, D42, and D49 to evaluate persistence of treatment effects. Animals were euthanized at study endpoint, and brains were collected 1 h following final treatment.

#### Experiment 3: Forelimb adjusting steps test crossover study

Experiment 3 evaluated whether sub-anesthetic ketamine alters parkinsonian motor impairment or interferes with the anti-parkinsonian effects of L-DOPA in female rats. For motor testing, female unilateral 6-OHDA-lesioned rats (*n* = 12) participated in a within-subject crossover experiment. The same animals underwent both ketamine and vehicle treatment conditions on separate treatment days. Sub-anesthetic ketamine (20 mg/kg, i.p.) or vehicle was administered using the 10-h treatment paradigm, and FAS performance was evaluated following the 1st, 4th, and 5th injections at 40-, 60-, and 80-min post-injection intervals. During the fifth injection condition, ketamine or vehicle treatment was paired with L-DOPA administration to determine whether ketamine altered L-DOPA-mediated motor benefit.

### L-DOPA-induced dyskinesia (LID) rating scale

LID severity was quantified by an investigator blinded to experimental condition using the well-established Abnormal Involuntary Movement (AIM) scale developed for the unilateral 6-OHDA rat model and validated across laboratories.^25,26^ AIMs were evaluated as limb, axial, and orolingual (LAO) behavioral subtypes. Limb AIMs consisted of purposeless movements of the forelimb contralateral to the lesion, axial AIMs consisted of dystonic posturing of the neck and upper body, and orolingual AIMs included vacuous jaw movements and tongue protrusions.

Behavioral assessments were conducted during 1-min observation periods performed every 20 min over 180 min following L-DOPA administration, consistent with established protocols.^4^ During each observation period, individual AIM subtypes were scored using a severity scale ranging from 0–4, where 0 represented absence of dyskinesia and 4 represented continuous, non-interruptible dyskinetic behavior. Total LAO-AIM scores were calculated by summing individual subtype scores across all time points. This behavioral approach is sensitive to pharmacological manipulation and correlates with molecular adaptations associated with LID.^10^ Comparison across AIM scoring methodologies has demonstrated strong validity and reproducibility within the unilateral 6-OHDA model.^26^

### Forelimb adjusting steps (FAS) test

The Forelimb Adjusting Steps (FAS) test was used to assess akinesia and forelimb motor function following unilateral dopamine depletion.^27,28^ Animals (n = 12) underwent FAS assessment before lesion surgery (Pre-LX) and following unilateral 6-OHDA lesion (Post-LX) to verify parkinsonian motor impairment. For treatment testing (Experiment 3), the same 12 animals participated in a within-subject crossover design in which ketamine and vehicle conditions were evaluated on separate treatment days. Sub-anesthetic ketamine (20 mg/kg, i.p.) or vehicle was administered using the 10-h dosing paradigm consisting of five injections delivered at 2-h intervals, with the fifth injection co-administered with L-DOPA.

Behavioral testing was conducted following the 1st, 4th, and 5th injections, and FAS performance was assessed at 40-, 60-, and 80-min post-injection time points. During testing, the hind limbs were elevated and one forepaw was restrained while the animal was moved laterally across a smooth surface (90 cm at 9 cm/s), such that body weight was supported by the unrestrained forelimb. Stepping behavior was evaluated using ipsilateral and contralateral forehand and backhand measures, which were combined to generate an overall stepping performance outcome. All behavioral sessions were video recorded and scored by a blinded investigator. Mean ± SEM stepping performance was calculated as previously described.^27,28^

### Tissue collection and Western blot analysis

Whole striatal tissue was collected from both hemispheres following euthanasia by CO₂ inhalation, with the right hemisphere designated as the lesioned side. Brains were rapidly removed, placed in a brain matrix, sectioned using razor blades, and striatal tissue was dissected on ice. Samples were flash-frozen and stored at −80°C until protein extraction. Tissue collection for Experiments 1 and 2 was performed 1 h following the final ketamine or vehicle injection. Striatal tissue was homogenized using a bead homogenizer in RIPA buffer supplemented with protease and phosphatase inhibitor mini tablets. Protein concentration was determined using the Pierce bicinchoninic acid (BCA) protein assay according to manufacturer instructions.

Protein samples were prepared at a concentration of 1 μg/μL, and 6.8 μg total protein was loaded per lane onto Any-kD Mini-PROTEAN TGX™ precast gels (Bio-Rad Laboratories, Hercules, CA). Proteins were transferred using a Trans-Blot Turbo Transfer System (Bio-Rad Laboratories). A Chameleon Duo prestained protein ladder (LI-COR, Lincoln, NE) was used for molecular weight determination. Membranes were blocked using LI-COR blocking buffer and incubated overnight at 4°C on a cold shaker with primary antibodies against tyrosine hydroxylase (TH; 1:2000; Millipore, Billerica, MA) and β-actin (1:2000; Sigma-Aldrich). Membranes were washed using TBST containing 0.1% Tween-20 and incubated for 1 h at room temperature with IRDye 800CW donkey anti-chicken secondary antibody (1:20,000; LI-COR) and IRDye 490CW donkey anti-goat secondary antibody (1:20,000; LI-COR). Blots were imaged using an Odyssey CLx imaging system (LI-COR) and analyzed using Image Studio Software (LI-COR). Protein expression was quantified using normalization to β-actin.

### Statistical Analyses

Statistical analyses were performed using GraphPad Prism version 10.6.1 software (GraphPad Software, Boston, MA, USA). For the attenuation study, total LAO-AIM scores were analyzed between vehicle-and ketamine-treated animals at D0, D7, and D14 using two-tailed Mann Whitney U tests with Holm Bonferroni correction for multiple comparisons across the three planned experimental days. LAO-AIM time-course data for D0, D7, and D14 were analyzed using mixed effects analysis with restricted maximum likelihood estimation, followed by Šídák’s multiple comparisons post hoc testing. For the established LID study, total LAO-AIM scores were analyzed between vehicle-and ketamine-treated animals at D0, D28, and D49 using two-tailed Mann–Whitney U tests with Holm–Bonferroni correction for multiple comparisons across the three planned experimental days. Total cumulative Area Under the Curve (AUC) scores across the 49-day study period were analyzed using an unpaired t-test. LAO-AIM time-course data at baseline, D0, D28, and D49 were analyzed using mixed-effects analysis with restricted maximum likelihood estimation, followed by Šídák’s multiple comparisons post hoc testing. Statistical significance was defined as *p* < 0.05, and data are presented as mean ± SEM. FAS data were analyzed using ANOVA models appropriate for the experimental comparison, followed by Tukey–Kramer corrected post hoc tests. Western blot quantification of TH/β-actin expression in intact versus lesioned striatal tissue was analyzed using a two-tailed paired *t* test. Time-course analyses and figure-specific statistical comparisons are described in the corresponding figure legends to ensure that each statistical test matches the structure of the data presented.

## Results

### Verification of lesion severity in Experiment 1: Female attenuation cohort

Prior to treatment assignment, unilateral nigrostriatal lesion severity was evaluated using amphetamine-induced rotational behavior, with FAS included as an additional behavioral measure of lesion-induced forelimb impairment. These behavioral assays are established markers of unilateral nigrostriatal dysfunction in 6-OHDA rodent models of PD.^4,27,28^ Following D-amphetamine administration (5 mg/kg, i.p.), animals that exhibited ≥4 net ipsilateral rotations/min were considered to have a severe unilateral lesion and were retained for subsequent analyses. This inclusion threshold is consistent with previous work from our laboratory showing that 6-OHDA injection into the medial forebrain bundle produces greater than 90% depletion of striatal dopamine content.

In the female attenuation cohort, amphetamine-induced rotational behavior was well above the inclusion criterion, with animals exhibiting 15.19 ± 1.96 net mean ± SEM ipsilateral rotations/min. These behavioral data supported successful induction of severe unilateral nigrostriatal lesions prior to treatment testing. At study completion, lesion severity was further confirmed by western blot analysis of striatal tissue. TH protein expression was normalized to β-actin, and normalized TH/β-actin values from the right lesioned striatum were compared with the matched left intact striatum within each animal. TH/β-actin expression was significantly reduced in the lesioned hemisphere relative to the intact hemisphere in both treatment groups (**Figure 1B–C**). Vehicle-treated animals showed a significant reduction in TH expression in the lesioned hemisphere compared with the intact hemisphere (paired two-tailed t test, t(9) = 15.36, *p* < 0.0001), and the same effect was observed in ketamine-treated animals (paired two-tailed t test, t(9) = 11.61, *p* < 0.0001). Representative western blot images showed reduced TH signal in lesioned striatal samples with preserved β-actin signal across lanes, supporting molecular verification of unilateral dopaminergic denervation in both treatment groups.

**Figure 1.**
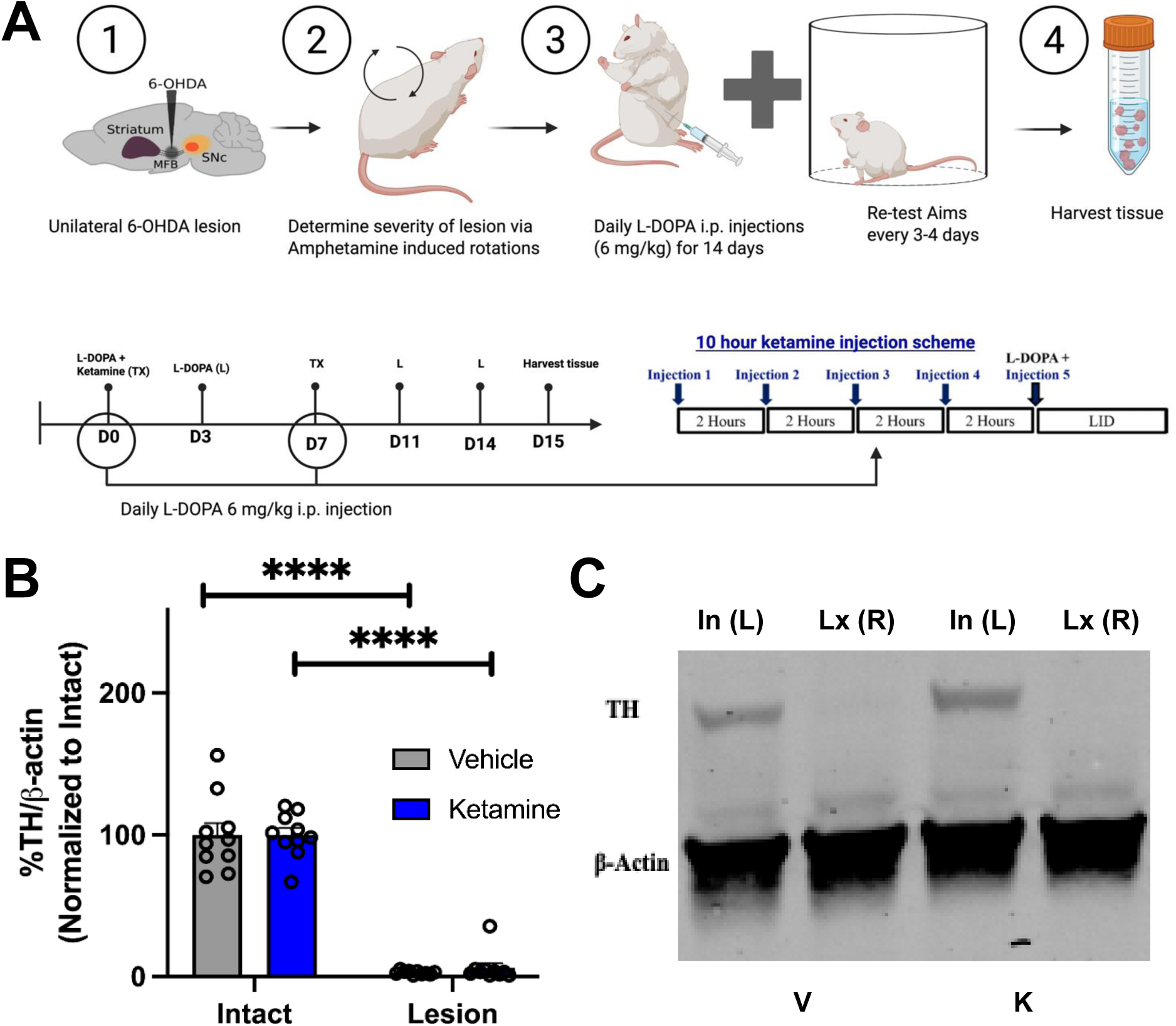
Study timeline and Verification of 6-ODHA lesion for Female Attenuation (Experiment 1). **(A)** Schematic overview of the experimental timeline. Adult female rats received a unilateral 6-OHDA lesion to induce a Parkinsonian phenotype, and lesion severity was verified using amphetamine-induced rotational testing. Animals were then administered daily L-DOPA (6 mg/kg, i.p.) for 14 days to induce dyskinesia. Sub-anesthetic ketamine (20 mg/kg, i.p.) or vehicle was delivered on treatment days D0 and D7 using a 10-hour paradigm consisting of five injections administered at 2-hour intervals; the fifth ketamine injection was co-administered with L-DOPA. Abnormal involuntary movements (AIMs) were assessed every 3–4 days throughout the study (n = 20). Animals were euthanized at the experimental endpoint for tissue collection. **(B)** Western blot quantification of TH protein expression normalized to β-actin confirmed successful dopaminergic denervation following unilateral 6-OHDA lesion. TH/β-actin levels were expressed as a percentage of the intact hemisphere. In both treatment groups, TH expression was significantly reduced in the lesioned hemisphere compared with the intact hemisphere, indicating robust lesion severity. In the vehicle group, TH expression was significantly decreased in the lesioned hemisphere compared with the intact hemisphere (paired two-tailed t-test, t(9) = 15.36, *p* < 0.0001). Similarly, in the ketamine group, TH expression was significantly decreased in the lesioned hemisphere compared with the intact hemisphere (paired two-tailed t-test, t(9) = 11.61, *p* < 0.0001). Data are presented as mean ± SEM; \*\*\*\**p* < 0.0001. **(C)** Representative western blot image showing TH protein expression in the intact left hemisphere [In (L)] and lesioned right hemisphere [Lx (R)] from vehicle-and ketamine-treated animals. Reduced TH signal in the lesioned hemisphere compared with the intact hemisphere visually confirms dopaminergic denervation in both treatment groups.

### Experiment 1: Ketamine attenuates L-DOPA-induced dyskinesia in female rats

To determine whether early ketamine administration attenuates the development of L-DOPA-induced dyskinesia (LID) in female rats, animals received repeated L-DOPA administration (6 mg/kg, i.p.) across a 14-day testing period following lesion verification and treatment assignment. Ketamine (20 mg/kg, i.p.) or vehicle was administered on Days 0 and 7 using the 10-h dosing paradigm consisting of five injections delivered every 2 h, with the fifth injection co-administered with L-DOPA. On non-treatment days, animals received L-DOPA alone, permitting assessment of dyskinesia severity in the absence of acute ketamine exposure and evaluation of sustained treatment effects.

We hypothesized based on the male data that we would see an acute effect of ketamine at D7 and also a long-term effect on D14, 1-week post-treatment. Vehicle-treated animals exhibited the expected progressive increase in total LAO-AIM scores across repeated L-DOPA exposure, reflecting gradual development of dyskinetic behavior. In contrast, ketamine-treated animals demonstrated attenuated dyskinesia severity across the experimental period (**Figure 2A**). Between-group comparisons were performed using two-tailed Mann–Whitney U tests with Holm–Bonferroni correction for multiple comparisons across D0, D7, and D14. No significant difference was detected between vehicle-and ketamine-treated animals on D0 (adjusted *p* = 0.0656). In contrast, ketamine significantly reduced LAO-AIM scores compared with vehicle on D7 (adjusted *p* = 0.0146) and D14 (adjusted *p* = 0.0015). Notably, D14 occurred one week following the final ketamine treatment session, indicating persistence of reduced dyskinesia severity beyond the acute treatment window. Together, these findings demonstrate that early ketamine administration altered the trajectory of LID development, resulting in sustained attenuation of dyskinetic behavior.

**Figure 2.**
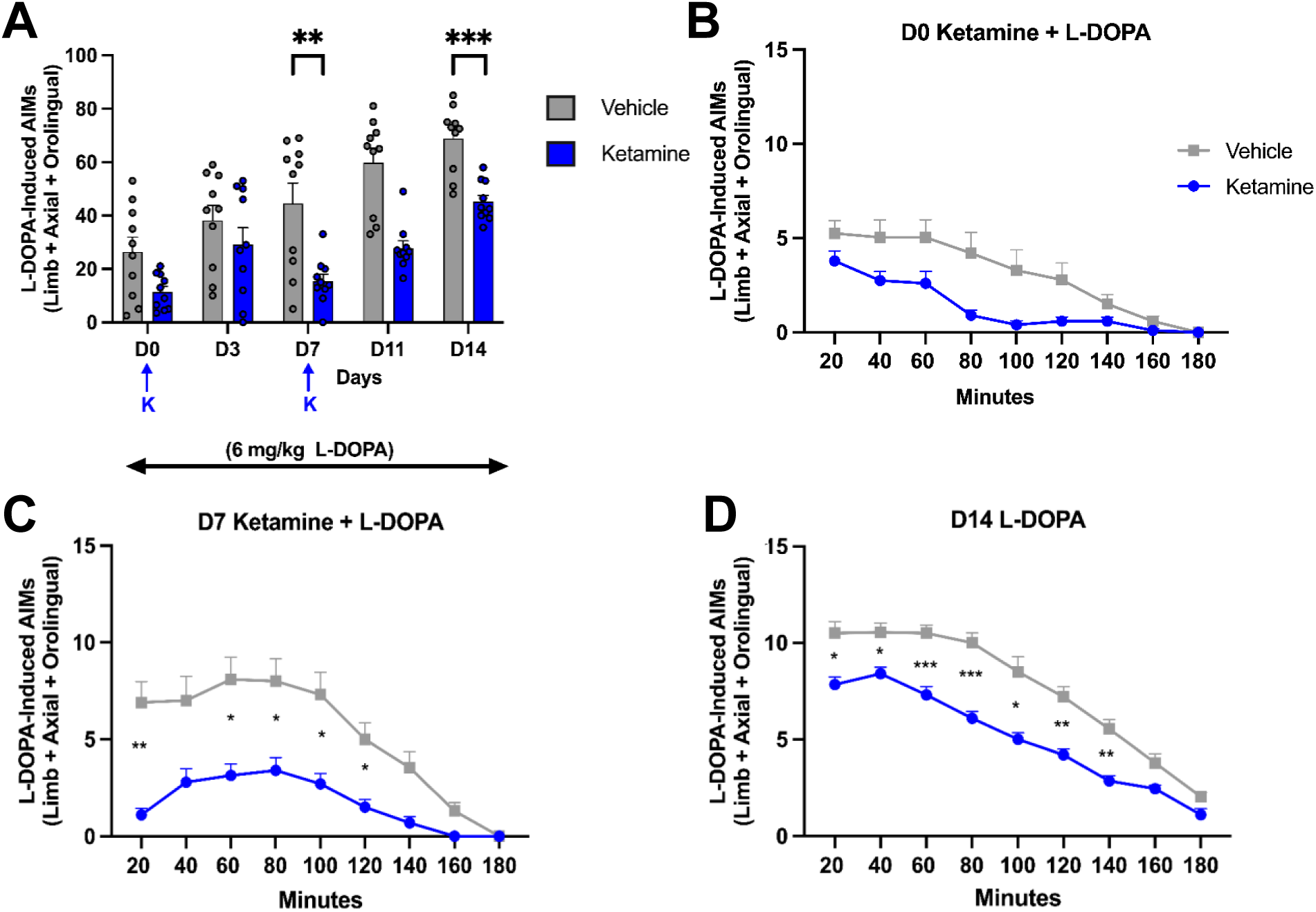
Sub-anesthetic ketamine attenuated the development of AIMs in the female rat model of LID. **(A)** Mean total LAO-AIM scores ± SEM across experimental time points. Female unilateral 6-OHDA-lesioned rats received daily L-DOPA administration (6 mg/kg, i.p.) and were treated with ketamine (20 mg/kg, i.p.; *n* = 10) or vehicle (*n* = 10) using the 10-h dosing paradigm on D0 and D7. Ketamine treatment attenuated the progressive development of dyskinesia, resulting in significantly reduced LAO-AIM severity on D7 and D14 compared with vehicle-treated animals. Blue arrows indicate treatment days. Between-group comparisons at D0, D7, and D14 were analyzed using two-tailed Mann–Whitney U tests. No significant difference was observed on D0 (*p* = 0.0656), whereas ketamine significantly reduced AIM scores on D7 (*p* = 0.0073) and D14 (***\*****p* = 0.0005). (B-D) Time-course analyses of LAO-AIM expression across the 180-min post-L-DOPA observation period. **(B)** D0 ketamine + L-DOPA, **(C)** D7 ketamine + L-DOPA, and **(D)** D14 L-DOPA alone. Ketamine-treated animals exhibited reduced dyskinesia severity relative to vehicle-treated animals across testing sessions, including D14, one week following the final ketamine treatment, when behavioral assessments were performed in the absence of acute ketamine exposure, indicating sustained attenuation of dyskinesia development. Time-course data were analyzed using mixed-effects models (REML) with Geisser–Greenhouse correction and appropriate multiple comparisons post hoc testing.

To further characterize dyskinesia expression throughout the behavioral session, LAO-AIMs were assessed every 20 min across a 180-min period following L-DOPA administration on Days 0, 7, and 14 (**Figure 2B–D**). On D0, corresponding to the initial treatment session, mixed-effects analysis (REML) revealed a significant main effect of time (*F*(2.607, 46.93) = 37.43, *p* < 0.0001), a significant main effect of treatment (*F*(1,18) = 6.172, *p* = 0.0230), and a significant time × treatment interaction (*F*(2.607, 46.93) = 4.519, *p* = 0.0098). These findings indicate reduced LAO-AIM expression in ketamine-treated animals across the behavioral session during the initial treatment day.

By D7, vehicle-treated animals demonstrated increased AIM severity across the behavioral session, consistent with progressive dyskinesia development, whereas ketamine-treated animals maintained reduced AIM expression throughout the time course. Mixed-effects analysis (REML) demonstrated a significant main effect of time (*F*(2.515,45.27) = 43.40, *p* < 0.0001), a significant main effect of treatment (*F*(1,18) = 16.18, *p* = 0.0008), and a significant time × treatment interaction (*F*(2.515,45.27) = 8.349, *p* = 0.0003), supporting robust separation between treatment groups during continued L-DOPA exposure.

Importantly, these treatment differences persisted on D14, when animals received L-DOPA alone in the absence of acute ketamine administration. Vehicle-treated animals exhibited pronounced dyskinesia across the behavioral session, whereas ketamine-treated animals continued to demonstrate reduced AIM expression (**Figure 2D**). Mixed-effects analysis revealed a significant main effect of time (*F*(4.251,76.51) = 230.4, *p* < 0.0001), a significant main effect of treatment (*F*(1,18) = 26.50, *p* < 0.0001), and a significant time × treatment interaction (*F*(4.251,76.51) = 6.608, *p* < 0.0001). Persistence of reduced dyskinesia severity one week following the final ketamine treatment suggests that ketamine attenuated LID in a sustained manner rather than producing only transient suppression of abnormal movements.

### Verification of lesion severity in Experiment 2: Established LID cohort

In the established LID cohort, amphetamine-induced rotational behavior also confirmed severe unilateral lesioning prior to experimental testing. Animals exhibited 14.41 ± 3.83 net mean ± SEM ipsilateral rotations/min, which was well above the ≥4 rotations/min inclusion threshold. Molecular confirmation of lesion severity was observed by western blot analysis at study completion. TH/β-actin expression was significantly reduced in the lesioned hemisphere relative to the intact hemisphere in both vehicle-and ketamine-treated animals (**Figure 3B–C**). In the vehicle group, TH expression was significantly decreased in the lesioned hemisphere compared with the intact hemisphere (paired two-tailed t test, t(4) = 3.393, *p* = 0.0275). Similarly, ketamine-treated animals showed a significant reduction in TH expression in the lesioned hemisphere relative to the intact hemisphere (paired two-tailed t test, t(4) = 3.623, *p* = 0.0223). Together, the amphetamine rotation data and TH western blot results confirmed successful unilateral dopaminergic denervation in the established LID cohort.

**Figure 3.**
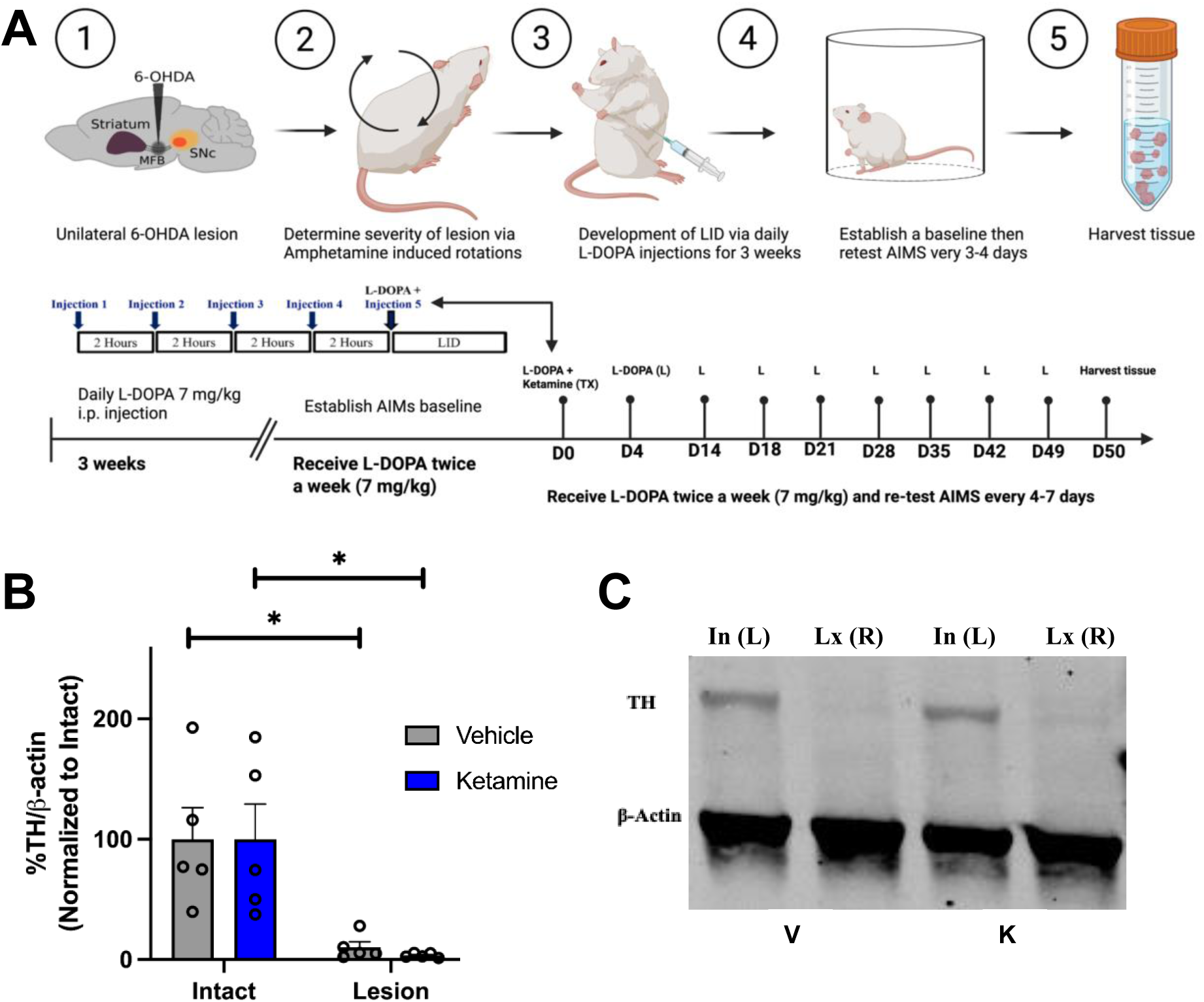
Study timeline and Verification of 6-ODHA lesion for Female Established (Experiment 2). **(A)** Schematic overview of the experimental timeline. Adult female rats received a unilateral 6-OHDA lesion to induce a Parkinsonian phenotype, and lesion severity was verified using amphetamine-induced rotational testing. Animals were then administered daily L-DOPA (7 mg/kg, i.p.) for 3 weeks induce dyskinesia. After 3 weeks a baseline of L-DOPA induced aims is established, and animals now receive maintenance L-DOPA (7 mg/kg, i.p.) injection twice a week. Sub-anesthetic ketamine (20 mg/kg, i.p.) or vehicle was delivered on treatment days D0 using a 10-hour paradigm consisting of five injections administered at 2-hour intervals; the fifth ketamine injection was co-administered with L-DOPA. Abnormal involuntary movements (AIMs) were assessed every 3–4 days throughout the study (N=10). Animals were euthanized at the experimental endpoint for tissue collection. **(B)** Western blot quantification of TH protein expression normalized to β-actin confirmed significant dopaminergic loss in the lesioned hemisphere compared with the intact hemisphere in both vehicle-treated animals (paired two-tailed t-test, t(4) = 3.393, *p* = 0.0275) and ketamine-treated animals (paired two-tailed t-test, t(4) = 3.623, *p* = 0.0223). TH/β-actin levels were expressed as a percentage of the intact hemisphere. Data are presented as mean ± SEM; \**p* < 0.05. **(C)** Representative western blot image showing reduced TH expression in the lesioned right hemisphere [Lx (R)] compared with the intact left hemisphere [In (L)] in vehicle-and ketamine-treated animals.

### Experiment 2: Ketamine reduces established L-DOPA-induced dyskinesia acutely and long-term in female rats

To determine whether ketamine reduces dyskinesia after abnormal involuntary movements were fully established, female rats received daily L-DOPA administration (7 mg/kg, i.p.) for three weeks to induce stable L-DOPA-induced dyskinesia (LID). AIMs were assessed throughout the induction period to verify development of robust dyskinetic behavior prior to treatment intervention. Following confirmation of established LID, animals were assigned to ketamine or vehicle treatment groups and underwent the 10-h ketamine treatment paradigm. Prior to treatment initiation, baseline AIM assessments (B1–B3) demonstrated comparable dyskinesia severity between groups, confirming equivalent behavioral status before intervention (**Figure 4A**). This ensured that subsequent differences in dyskinesia expression reflected treatment effects rather than pre-existing variability in AIM severity.

**Figure 4.**
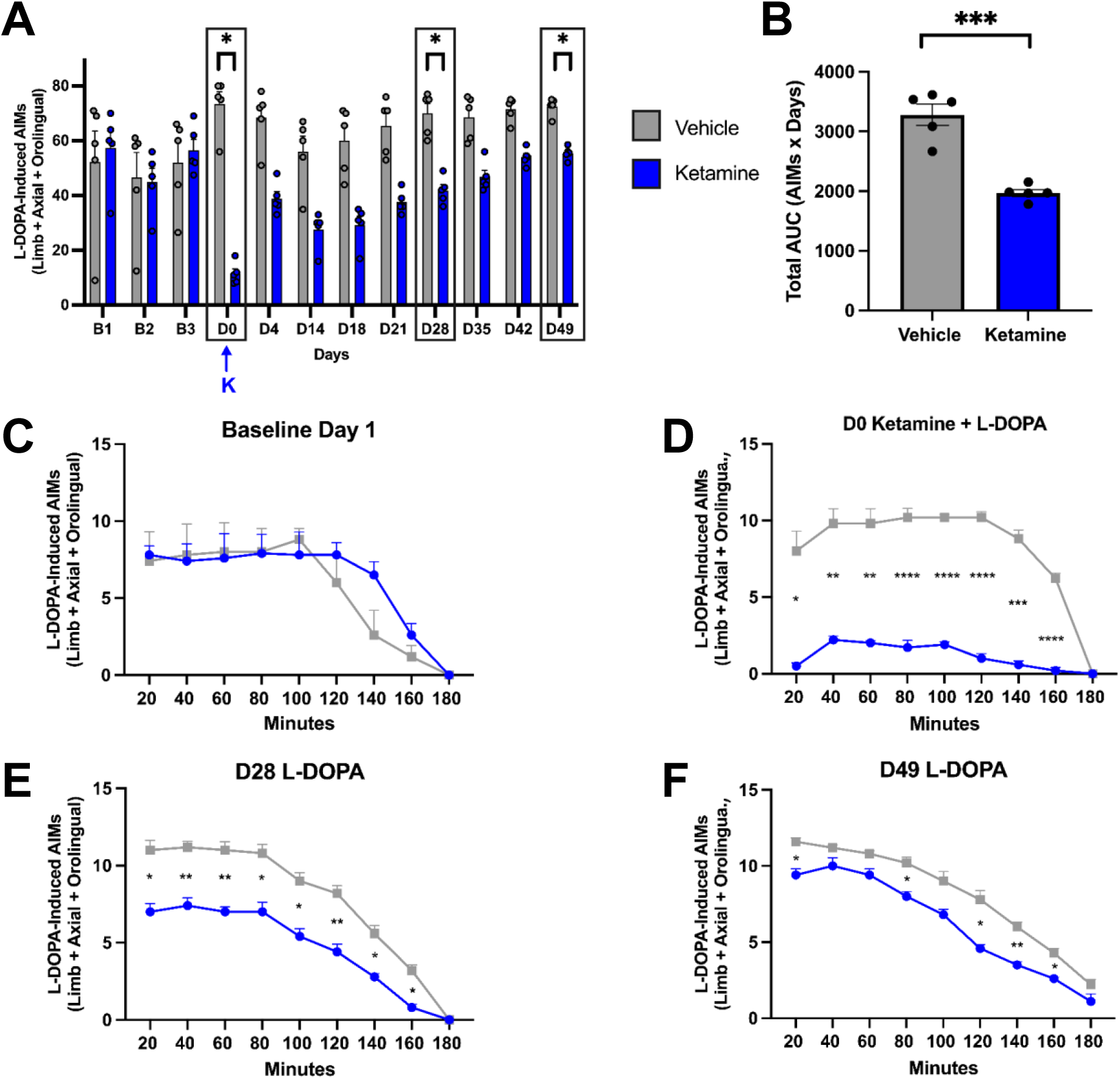
Sub-anesthetic ketamine reduced established L-DOPA-induced dyskinesia in the female rat model of LID. **(A)** Total limb, axial, and orolingual abnormal involuntary movement (LAO AIM) scores across testing days in unilateral 6-OHDA-lesioned rats. Animals received vehicle (n = 5) or ketamine (20 mg/kg, i.p.; n = 5) via a 10-h dosing paradigm on Day 0 (blue arrow). Ketamine significantly reduced dyskinesia compared to vehicle on D0, D28, and D49 (two-tailed Mann–Whitney U tests with Holm correction; U = 0, Holm-adjusted \**p* = 0.0238 for each day). **(B)** Total cumulative AIMs scores calculated via Area Under the Curve (AUC) analysis for each individual animal across the entire 49-day study period. A single day of ketamine treatment significantly suppressed overall L-DOPA-induced AIMs across the duration of the study (unpaired t-test; t(8) = 6.958, \*\*\**p* = 0.0001, R^2^ = 0.8582). **(C–F)** Time-course analyses of LAO-AIM expression across the 180-min post-L-DOPA observation period at key experimental time points. **(C)** Baseline Day 1 demonstrating comparable dyskinesia profiles between groups prior to treatment. **(D)** D0 Ketamine + L-DOPA demonstrating marked acute suppression of established dyskinesia following the 10-h ketamine treatment paradigm. **(E)** D28 L-DOPA showing sustained reductions in LAO-AIM expression in ketamine-treated animals relative to vehicle controls. **(F)** D49 L-DOPA demonstrating persistence of the anti-dyskinetic effect at later experimental time points, although significant attenuation was limited to selected portions of the behavioral session. Time-course data were analyzed using mixed-effects models (REML) with Geisser–Greenhouse correction followed by Šídák multiple comparisons post hoc tests.

We hypothesized based on the male data that we would see an acute effect of ketamine at D0 and also long-term effects, 1-month post-treatment on D28 and at the end of the Experiment on D49. On treatment day D0, ketamine-treated female rats showed a marked reduction in total LAO AIM scores compared with vehicle-treated controls (**Figure 4A**). Vehicle-and ketamine-treated animals were compared on D0, D28, and D49 using nonparametric two-tailed Mann–Whitney U tests with Holm-Bonferroni correction for multiple comparisons. Ketamine significantly reduced total LAO AIM scores compared with vehicle on D0, D28, and D49 (U = 0 for all comparisons; Holm-Bonferroni-adjusted *p* = 0.0238 for each comparison). Although the adjusted p-values were identical because each comparison showed complete rank separation between vehicle-and ketamine-treated animals, the largest reduction was observed on D0, with smaller but significant reductions persisting through D28 and D49. Together, these findings indicate that sub-anesthetic ketamine reduced established LID in female rats both acutely and at later experimental time points.

To evaluate the cumulative movement severity over the entire 49-day study period, total Area Under the Curve (AUC) scores were calculated for each individual animal. An unpaired t-test on these values confirmed a highly significant reduction in total AIMs scores in the ketamine-treated group (n = 5) compared to vehicle controls (n = 5) (Vehicle AUC: 3279 ± 119.8; Ketamine AUC: 1971 ± 66.73; t(8) = 6.958, *p* = 0.0001, R^2^ = 0.8582; **Figure 4B**). These cumulative data underscore a robust and lasting overall suppression of L-DOPA-induced dyskinesia following a single day of ketamine treatment.

To further characterize the temporal dynamics of dyskinesia expression, LAO-AIMs were analyzed across the 180-min post-L-DOPA testing session at baseline and following ketamine treatment (**Figure 4C–F**). At baseline (B1), ketamine-assigned and vehicle-assigned animals exhibited comparable AIM profiles across the behavioral session, confirming similar magnitudes and temporal patterns of dyskinesia prior to treatment intervention (**Figure 4C**). Immediately following the 10-h treatment paradigm (D0), ketamine markedly attenuated established dyskinesia across the behavioral session (**Figure 4D**). Šídák post hoc comparisons following mixed-effects analysis (REML) demonstrated significantly reduced AIM scores in ketamine-treated animals relative to vehicle controls at 20, 40, 60, 80, 100, 120, 140, and 160 min following L-DOPA administration (*p* = 0.0355, *p* = 0.0085, *p* = 0.0098, *p* < 0.0001, *p* < 0.0001, *p* < 0.0001, *p* = 0.0003, and *p* < 0.0001, respectively). Together, these findings indicate robust acute suppression of established dyskinesia across both peak and non-peak phases of the behavioral session.

To evaluate durability of this effect, dyskinesia expression was reassessed during continued L-DOPA exposure at later experimental time points. At D28, mixed-effects analysis (REML) revealed a significant main effect of time (*F*(3.219,25.75) = 199.8, *p* < 0.0001), a significant main effect of treatment (*F*(1,8) = 46.40, *p* = 0.0001), and a significant time × treatment interaction (*F*(3.219,25.75) = 7.209, *p* = 0.0009). Ketamine-treated animals maintained reduced LAO-AIM expression across the 180-min testing session relative to vehicle-treated controls, indicating sustained attenuation of dyskinetic behavior during ongoing L-DOPA administration (**Figure 4E**).

This effect remained detectable through D49, although treatment differences became more temporally restricted across the behavioral session (**Figure 4F**). Šídák post hoc comparisons demonstrated significantly lower AIM scores in ketamine-treated animals at 20, 40, 100, 120, 140, and 160 min following L-DOPA administration (*p* = 0.0009, *p* = 0.0191, *p* = 0.0009, *p* = 0.0009, *p* < 0.0001, and *p* = 0.0001, respectively). No significant differences were detected at 60, 80, or 180 min. These findings indicate persistence of ketamine’s anti-dyskinetic effects at later experimental stages, although attenuation was no longer uniform across the entire behavioral session.

Collectively, these results demonstrate that sub-anesthetic ketamine reduces established L-DOPA-induced dyskinesia in female rats across both short-and long-term timescales. The observed behavioral profile includes immediate suppression of established dyskinesia following treatment administration as well as sustained attenuation during continued L-DOPA exposure, supporting prolonged modulation of dyskinesia expression following ketamine treatment.

### Verification of lesion severity in Experiment 3: Female FAS cohort

In the female FAS cohort, amphetamine-induced rotational behavior confirmed severe unilateral lesioning, with animals exhibiting 13.97 ± 1.82 net mean ± SEM ipsilateral rotations/min prior to behavioral testing. FAS was then used as an additional behavioral measure of lesion-induced forelimb impairment and motor response to treatment. Western blot analysis provided additional molecular confirmation of unilateral dopaminergic denervation in this cohort (**Figure 5B–C**). Across paired intact and lesioned striatal samples, TH/β-actin expression was significantly reduced in lesioned tissue compared with intact tissue (paired two-tailed t test, t(11) = 2.236, *p* = 0.0470, n = 12 pairs). Representative western blot images demonstrated reduced TH signal in lesioned striatal samples relative to matched intact samples from the same FAS crossover cohort, in which animals received both vehicle and ketamine on separate testing days. Collectively, the behavioral lesion assessments and western blot findings confirmed severe unilateral nigrostriatal lesions across all three experimental cohorts.

**Figure 5.**
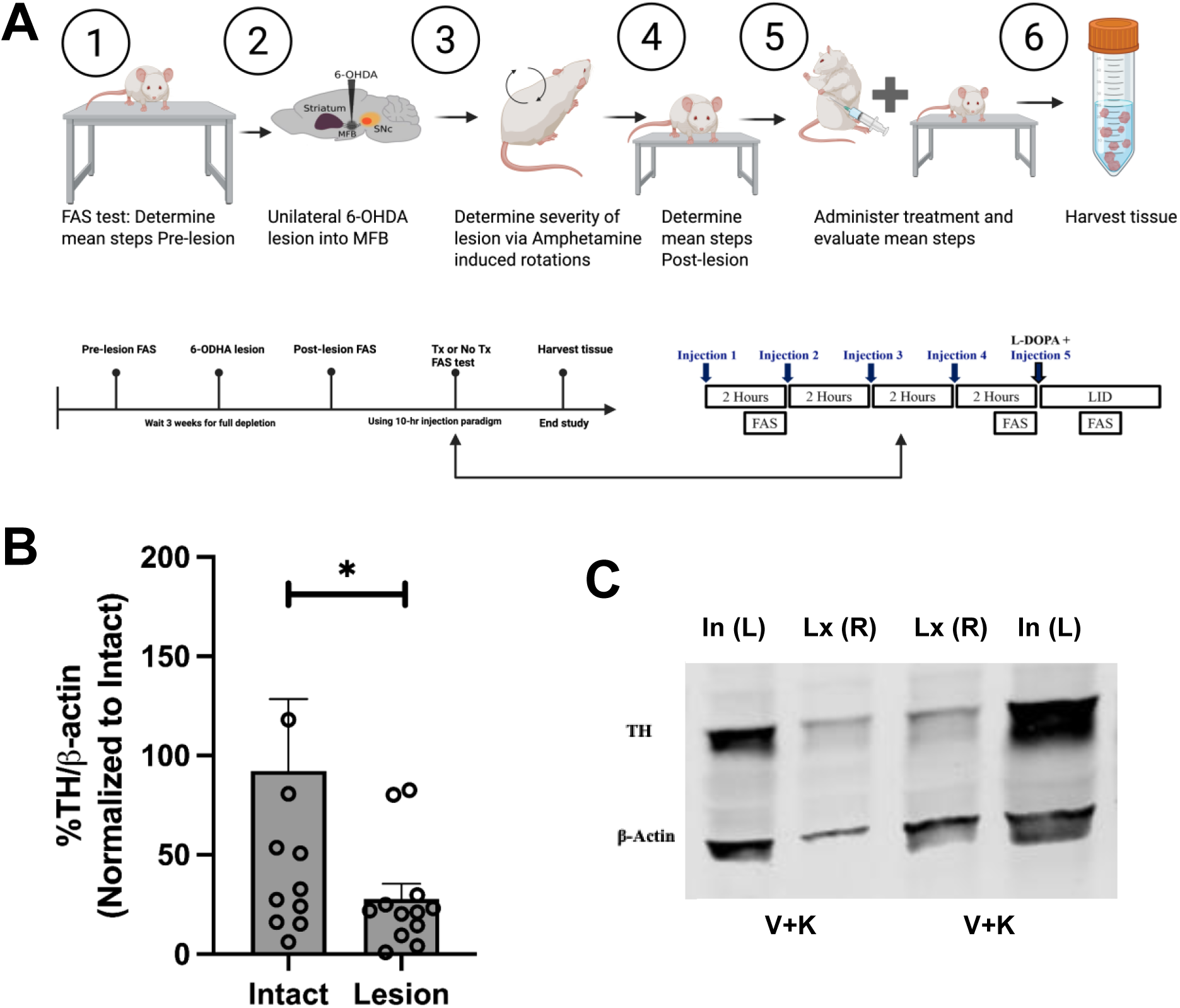
Study timeline and Verification of 6-ODHA lesion for Female FAS (Experiment 3). **(A)** Schematic overview of the FAS experimental design. Adult female rats were tested before 6-OHDA lesion surgery (Pre-LX; *N* = 12), lesioned unilaterally, and verified by amphetamine-induced rotational testing. FAS was repeated after lesion verification (Post-LX; *N* = 12). The same animals then completed a within-subject crossover study in which ketamine (20 mg/kg, i.p.) or vehicle was administered on separate testing days using a 10-h, five-injection paradigm, with the fifth injection co-administered with L-DOPA. FAS was assessed after the 1st, 4th, and 5th injections by a blinded investigator. The crossover analysis included 24 treatment sessions total (*N* = 12 ketamine; *N* = 12 vehicle). Animals were euthanized at the experimental endpoint for tissue collection. **(B)** Quantification of tyrosine hydroxylase (TH) protein expression normalized to β-actin in intact and lesioned striatal tissue. TH/β-actin levels were significantly reduced in the lesioned striatum compared with the intact striatum, confirming successful unilateral dopaminergic lesioning. Data are expressed as percent of the intact side and presented as mean ± SEM with individual animals shown. Statistical analysis was performed using a two-tailed paired *t* test; *t*(11) = 2.236, *p* = 0.0470, *N* = 12 pairs. *p* < 0.05. **(C)** Representative Western blot images showing TH and β-actin bands from intact (In) and lesioned (Lx) striatal samples. These samples were collected from the crossover FAS cohort, in which the same animals received both vehicle/saline and ketamine on separate treatment days. Reduced TH signal in the lesioned striatum relative to the intact striatum further verifies the presence of a unilateral 6-OHDA lesion.

### Experiment 3: Ketamine does not interfere with L-DOPA efficacy and exhibits acute anti-parkinsonian activity in female parkinsonian rats

In Experiment 3, we evaluated whether ketamine alters the anti-parkinsonian efficacy of L-DOPA or independently influences motor performance in female rats. Because an effective anti-dyskinetic therapy should reduce AIMs without compromising the therapeutic motor benefits of dopaminergic treatment, motor performance was assessed using the FAS test in unilateral 6-OHDA-lesioned female rats using a within-subject crossover design. Successful lesion induction was confirmed behaviorally prior to treatment assessment. Following unilateral 6-OHDA lesioning, animals exhibited a marked reduction in stepping performance relative to pre-lesion baseline, confirming the establishment of parkinsonian motor impairment. Median stepping performance decreased from 99.71 before lesioning to 49.45 after lesioning (Mann–Whitney U = 0, exact two-tailed *p* < 0.0001, n = 12 per condition; (**Figure 6A**).

**Figure 6.**
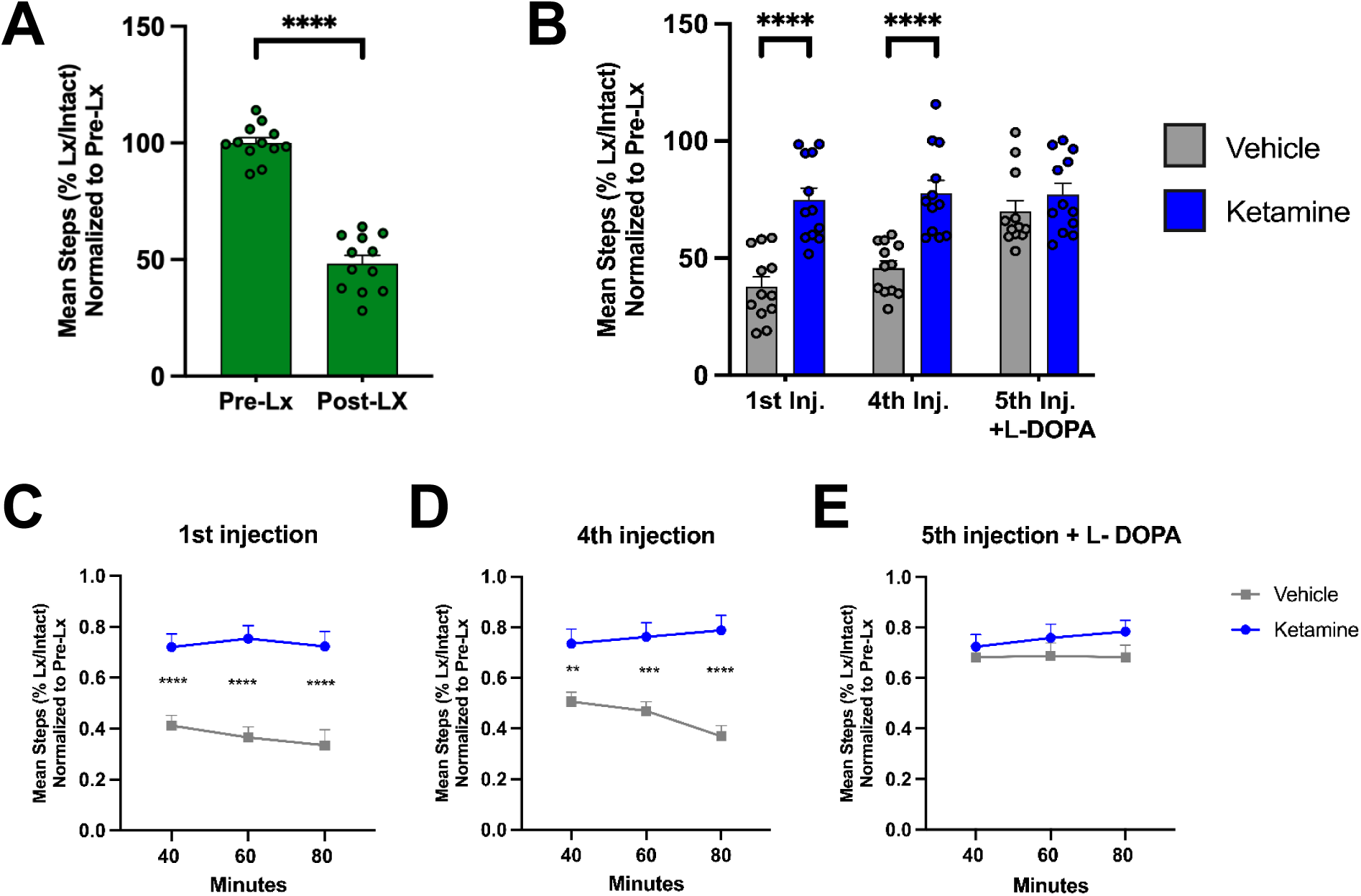
Ketamine did not interfere with L-DOPA efficacy and exhibited acute anti-parkinsonian activity in female Parkinsonian rats. **(A)** Forelimb adjusting steps (FAS) performance before (Pre-Lx) and after (Post-Lx) unilateral 6-OHDA lesion. Female rats exhibited a significant reduction in stepping performance following lesion induction, confirming successful establishment of parkinsonian motor impairment. Data are presented as mean ± SEM (*n* = 12). **(B)** Quantification of FAS performance across treatment injections. Using a within-subject crossover design, the same animals received ketamine (20 mg/kg, i.p.; blue) and vehicle (grey) treatment conditions on separate experimental days. Ketamine-treated animals exhibited significantly improved stepping performance relative to vehicle during the first and fourth injections. During the fifth injection, when ketamine or vehicle was co-administered with L-DOPA, ketamine did not impair L-DOPA-mediated motor improvement. Data are presented as mean ± SEM (*n* = 12). Statistical analysis was performed using one-way ANOVA followed by Tukey–Kramer multiple comparisons post hoc tests (*F*(5,66) = 13.14, \*\*\*\**p* < 0.0001). **(C–E)** Time-course analyses of FAS performance measured at 40, 60, and 80 min post-injection. **(C)** First injection and **(D)** fourth injection, during which ketamine significantly improved stepping performance relative to vehicle across all time points. **(E)** Fifth injection following co-administration with L-DOPA, showing no significant difference between vehicle + L-DOPA and ketamine + L-DOPA conditions. Time-course data were analyzed using mixed-effects models (REML) followed by Šídák multiple comparisons post hoc tests.

We hypothesized based on the male data that we would see an acute anti-parkinsonian effect of ketamine, and that ketamine would not interfere with the therapeutic action of L-DOPA. To determine whether ketamine independently affects motor performance, animals underwent vehicle and ketamine treatment conditions on separate experimental days, and FAS performance was evaluated following the first and fourth injections of the 10-h treatment paradigm. Quantification across treatment injections demonstrated significantly improved stepping performance during ketamine treatment conditions relative to vehicle controls (one-way ANOVA with Tukey–Kramer multiple comparisons post hoc testing, *F*(5,66) = 13.14, \*\*\*\**p* < 0.0001; **Figure 6B**).

Temporal analysis of FAS performance further characterized these behavioral effects (**Figure 6C–E**). Following the first injection, mixed-effects analysis (REML) with Šídák multiple comparisons post hoc testing demonstrated significantly improved stepping performance during ketamine treatment relative to vehicle treatment at 40, 60, and 80 min post-injection (all *p* < 0.0001; **Figure 6C**). Similarly, following the fourth injection, ketamine-treated animals continued to exhibit significantly greater stepping performance at 40 min (*p* = 0.0047), 60 min (*p* = 0.0003), and 80 min (*p* < 0.0001) relative to vehicle treatment (**Figure 6D**). Because these assessments occurred prior to L-DOPA co-administration, these findings indicate that ketamine was associated with improved stepping performance independent of acute dopaminergic treatment.

To directly evaluate whether ketamine interferes with the therapeutic motor effects of L-DOPA, FAS performance was assessed following the fifth injection, during which ketamine or vehicle treatment was paired with L-DOPA administration. As expected, L-DOPA improved stepping performance under both treatment conditions. Importantly, no significant differences were detected between vehicle + L-DOPA and ketamine + L-DOPA treatment conditions at 40 min (*p* = 0.8902), 60 min (*p* = 0.6160), or 80 min (*p* = 0.3338) following administration (**Figure 6E**). These findings indicate that ketamine did not impair the anti-parkinsonian efficacy of L-DOPA. Collectively, these findings demonstrate that sub-anesthetic ketamine exhibits acute anti-parkinsonian activity in female 6-OHDA-lesioned rats while preserving responsiveness to dopaminergic therapy. Ketamine treatment improved motor performance during standalone treatment conditions and remained compatible with L-DOPA-mediated restoration of motor function.

## Discussion

The present study demonstrates that sub-anesthetic ketamine exerts anti-dyskinetic and anti-parkinsonian effects in female rat models of L-DOPA-induced dyskinesia (LID). Ketamine attenuated the development of dyskinesia when administered during early L-DOPA exposure and reduced the severity of established dyskinesia after abnormal involuntary movements (AIMs) had developed. These effects persisted beyond the acute treatment period, indicating that ketamine produces sustained modulation of dyskinesia rather than transient behavioral suppression. Ketamine also improved forelimb stepping under standalone treatment conditions and did not compromise the therapeutic motor benefit of L-DOPA. Collectively, these findings extend previous observations in male models and demonstrate that the anti-dyskinetic and anti-parkinsonian effects of sub-anesthetic ketamine are also present in female preclinical models.^4,19^

### Glutamatergic signaling, synaptic plasticity, and sustained ketamine effects

The anti-dyskinetic effects observed in the present study are consistent with extensive evidence implicating dysregulated glutamatergic transmission and N-methyl-D-aspartate (NMDA) receptor signaling in LID pathophysiology. Chronic L-DOPA treatment promotes maladaptive corticostriatal plasticity characterized by altered dopamine-glutamate interactions, receptor regulation, and synaptic homeostasis, ultimately contributing to sensitization of striatal circuitry and pathological motor output.^3,10^ At the synaptic level, Picconi et al. demonstrated that LID is associated with impaired bidirectional striatal plasticity, including abnormalities in both long-term potentiation and long-term depression.^11^ Gardoni et al. further identified alterations in NMDA receptor composition and receptor-scaffolding interactions within dyskinetic striatal circuitry.^12^ Together, these findings implicate abnormal glutamatergic signaling and NMDA receptor-dependent plasticity as important components of LID and provide a mechanistic rationale for targeting these pathways therapeutically.

The clinical efficacy of amantadine further supports NMDA receptor modulation as a therapeutic strategy for LID. Amantadine reduces dyskinesia severity and remains an important pharmacological option for patients with LID.^3,25^ However, its clinical utility can be limited by neuropsychiatric adverse effects, incomplete therapeutic responses, and variability in the durability of benefit.^10,25^ The sustained effects observed with ketamine may therefore be particularly relevant. Although both ketamine and amantadine influence NMDA receptor signaling, ketamine engages additional downstream processes involved in glutamatergic transmission and activity-dependent plasticity.^13,15,17,18^ These broader neurobiological actions may contribute to the prolonged behavioral effects observed following sub-anesthetic ketamine treatment.

A major finding of the present study was that reductions in dyskinesia persisted after the acute ketamine treatment period. Ketamine is capable of rapidly altering glutamatergic signaling while also engaging longer-lasting molecular pathways associated with synaptic remodeling.^13,17,18^ In neuropsychiatric models, NMDA receptor blockade has been linked to activity-dependent BDNF signaling, enhanced AMPA receptor-mediated transmission, and downstream pathways associated with synaptogenesis and structural plasticity.^15,16,17,18^ Although these mechanisms have been characterized primarily in the context of depression, they provide a potential framework for understanding how a relatively brief ketamine exposure could produce behavioral effects that persist beyond its acute pharmacological action.

Within the context of LID, ketamine may therefore do more than transiently suppress dyskinetic output. Its actions on glutamatergic signaling and plasticity-related pathways may promote longer-lasting changes within corticostriatal networks that have undergone maladaptive reorganization during chronic L-DOPA treatment.^3,10^ This interpretation is consistent with the sustained reduction in AIM scores observed in the present study and with previous work demonstrating prolonged anti-dyskinetic effects of sub-anesthetic ketamine in male models.^4,19,20,31,32^ Importantly, clinical observations also provide preliminary support for sustained effects: case reports have described prolonged reductions in LID following sub-anesthetic ketamine infusion, and a Phase I open-label study has evaluated low-dose ketamine infusion as a potential treatment for LID in PD.^33,34^ Although these clinical observations remain limited, they support continued investigation of ketamine as a treatment capable of producing effects beyond the period of acute drug exposure.

### Biological sex and ketamine responsiveness

An important contribution of the present study is the demonstration that ketamine retains therapeutic efficacy in female models of PD and LID. This is particularly relevant given the historical underrepresentation of females in preclinical neuroscience and pharmacological research.^1,2^ Biological sex can influence dopaminergic signaling, glutamatergic transmission, and synaptic plasticity, all of which are relevant to both dyskinesia pathophysiology and ketamine pharmacology.^5,6,8^ Estrogens can modulate striatal dopamine synthesis, release, receptor function, and neuronal responsiveness, providing a biological basis by which female neurobiology could influence both LID development and pharmacological response.^8,9^

Evidence from preclinical models of depression further demonstrates that ketamine responsiveness can vary according to sex and ovarian hormone status. Carrier and Kabbaj reported greater sensitivity to low-dose ketamine in female rats compared with males, with this enhanced behavioral response dependent in part on ovarian hormones.^21^ Dossat et al. similarly demonstrated sex-and estrous-related differences in behavioral and biochemical responses to low-dose ketamine.^22^ In addition to these pharmacodynamic differences, Saland and Kabbaj reported sex-dependent ketamine pharmacokinetics, including differences in plasma and brain exposure to ketamine and its metabolites following comparable dosing.^23^ Together, these findings indicate that sex can influence multiple determinants of ketamine responsiveness and reinforce the importance of establishing therapeutic efficacy directly in female models rather than extrapolating exclusively from male studies.

Despite these potential sources of biological variability, ketamine produced clear anti-dyskinetic and anti-parkinsonian effects in the female cohorts examined here. The present study was not designed to determine whether ketamine efficacy differs quantitatively between males and females or to define the contribution of ovarian hormones to treatment response. Rather, the primary objective was to establish whether effects previously demonstrated in male LID models are also observed in females.^4,19^ The present findings address this question and provide a foundation for future studies directly examining sex-dependent determinants of ketamine efficacy.

### Anti-parkinsonian effects and preservation of L-DOPA motor benefit

An essential consideration for any candidate LID therapy is whether dyskinesia can be reduced without compromising the therapeutic effects of L-DOPA. L-DOPA remains the most effective pharmacological treatment for parkinsonian motor symptoms, despite the emergence of dyskinesia with chronic treatment.^35^ Any adjunctive therapy that reduces AIMs at the expense of L-DOPA-mediated motor improvement would therefore have limited clinical utility.

The forepaw adjusting steps (FAS) experiment provided two important findings. First, following unilateral 6-OHDA lesion, animals displayed substantial forelimb impairment, consistent with parkinsonian motor dysfunction (**Figure 6A**). During standalone treatment conditions, ketamine improved stepping performance relative to vehicle during both the first and fourth injections of the 10-h dosing paradigm (**Figure 6B-D**). Because these assessments occurred before L-DOPA co-administration, the improvement suggests an acute anti-parkinsonian effect of ketamine that was independent of immediate dopaminergic replacement.

Second, ketamine did not interfere with the therapeutic motor effects of L-DOPA. Following the fifth injection, when ketamine or vehicle was administered in combination with L-DOPA, stepping performance improved under both conditions, with no significant impairment in animals receiving ketamine + L-DOPA relative to vehicle + L-DOPA (**Figure 6E**). Thus, ketamine reduced dyskinetic behavior while preserving L-DOPA-mediated improvement in forelimb motor function.

These findings argue against generalized motor suppression as an explanation for ketamine’s anti-dyskinetic effects. Instead, ketamine appears capable of reducing pathological dyskinetic output while maintaining, and under standalone conditions improving, motor performance. This distinction is mechanistically relevant because L-DOPA restores dopamine-dependent signaling required for movement but, with repeated exposure, also drives maladaptive plasticity that contributes to LID.^3,25^ Ketamine may therefore preferentially modulate non-dopaminergic processes associated with pathological plasticity without preventing the dopamine-dependent signaling required for therapeutic motor benefit.

Recent electrophysiological findings provide a circuit-level framework for this interpretation. LID is associated with abnormal patterns of cortical and corticostriatal activity, including dyskinesia-related oscillatory activity, and sub-anesthetic ketamine has been shown to suppress specific LID-associated spectral abnormalities.^31^ More recent work further demonstrated that dyskinesia is accompanied by disrupted relationships between motor cortical activity and movement and that ketamine can reorganize abnormal neuronal interactions within motor cortical networks.^36,37^ These findings suggest that ketamine may reduce dyskinesia by modifying pathological network dynamics rather than broadly suppressing motor-system activity, which could help explain the coexistence of anti-dyskinetic activity and preserved L-DOPA-mediated motor improvement.

NMDA receptor modulation is unlikely to account for all of ketamine’s behavioral effects. Ketamine and several of its metabolites have also been reported to allosterically modulate opioid receptors,^14^ and previous work from our laboratory demonstrated that opioid receptor antagonism differentially influences the anti-dyskinetic and anti-parkinsonian effects of sub-anesthetic ketamine.^19^ These findings raise the possibility that ketamine’s therapeutic profile in LID reflects coordinated modulation of multiple signaling systems, including glutamatergic and opioid receptor-associated mechanisms. Further work will be required to determine how these pathways interact and whether they contribute differentially to the acute and sustained behavioral effects of ketamine.

Interpretation of the FAS findings should nevertheless remain appropriately constrained. The FAS assay primarily measures forelimb akinesia and does not capture the full spectrum of parkinsonian motor dysfunction.^26,28^ Additional behavioral measures, including spontaneous forelimb use, skilled reaching, gait analysis, balance and coordination tasks, and rotational behavior, could provide a more comprehensive assessment of ketamine’s motor effects.^26,38,39^ Thus, although the present findings support an anti-parkinsonian effect of ketamine in the FAS assay, additional studies are necessary to determine the breadth of this motor benefit.

## Limitations and future directions

Several limitations should be considered when interpreting the present findings. First, although the study establishes ketamine efficacy in female animals, it was not designed as a direct comparison of males and females. The results therefore demonstrate that ketamine’s anti-dyskinetic and anti-parkinsonian effects are present in females but do not establish the magnitude or nature of sex differences in treatment response.

Second, estrous cycle stage was not monitored. Ovarian hormones are known to influence dopaminergic signaling and ketamine responsiveness,^8,21,22,23^ and hormonal state could therefore contribute to interindividual variability in behavioral effects. Systematic assessment of estrous stage, ovariectomy, or hormone replacement will be important for determining whether ovarian hormone fluctuations modify ketamine efficacy in LID. However, the absence of estrous staging does not detract from the primary objective of the present study, which was to determine whether ketamine retains efficacy in female animals under conditions that incorporate naturally occurring biological variability.

Third, the present experiments were primarily behavioral and were not designed to identify the molecular or circuit mechanisms responsible for the sustained effects of ketamine. The existing literature implicates glutamatergic signaling, plasticity-related pathways, cortical network activity, and potentially opioid receptor signaling, but these mechanisms were not directly tested here.^14,15,17,18,31,36,37^ Future studies combining behavioral analysis with electrophysiological, molecular, and circuit-level approaches will be necessary to determine which mechanisms are required for ketamine’s acute and sustained therapeutic effects.

Finally, although the unilateral 6-OHDA model is well established for studying LID, no preclinical model fully reproduces the complexity of human PD.^25^ Nevertheless, the present findings add to converging evidence from male preclinical studies,^4,19,20^ clinical case reports,^33^ and early clinical investigation^34^ supporting continued evaluation of sub-anesthetic ketamine for dyskinesia. Demonstrating efficacy in female animals further strengthens the translational foundation for this work by establishing that the therapeutic effects are not restricted to male preclinical models.

## Conclusion

The present study demonstrates that sub-anesthetic ketamine attenuates both the development and established expression of L-DOPA-induced dyskinesia in female rats while also improving parkinsonian motor performance and preserving the therapeutic motor effects of L-DOPA. The persistence of anti-dyskinetic effects beyond the acute treatment period is consistent with the possibility that ketamine engages longer-lasting plasticity-and circuit-related mechanisms in addition to acute NMDA receptor modulation. Importantly, these findings extend previous work in male models and demonstrate that ketamine’s anti-dyskinetic and anti-parkinsonian effects are also present in females.^4,19^ Together, these results strengthen the preclinical rationale for continued investigation of sub-anesthetic ketamine as a potential adjunctive therapy for LID and provide a foundation for future studies examining the molecular, circuit, and sex-dependent determinants of treatment response.

## CRediT authorship contribution statement

**Raveena Parmar**: Formal analysis, Investigation, Methodology, Validation, Writing – original draft. **Mitchell J. Bartlett**: Formal analysis, Investigation, Methodology, Supervision, Validation, Resources, Writing – review & editing. **Jasmine R. Meredith**: Investigation, Methodology. **Sansita Singh**: Investigation, Methodology. **Sydney Ragsdale**: Investigation, Methodology. **Scott J. Sherman**: Writing – review & editing. **Torsten Falk**: Conceptualization, Funding acquisition, Resources, Supervision, Writing – review & editing.

## Acknowledgements

Research support: National Institute of Health NINDS [R01 NS122805, TF], Training grant funding NIH T35 HL007479 [MJB], NIH NINDS Diversity Supplement R01 NS122805-03S1 [RP], Robert & Payton Davies PD Research Fund and Mary Mack Estate Gift (SJS, TF).

